# Partially redundant actin genes in *Chlamydomonas* control flagellum-directed traffic and transition zone organization

**DOI:** 10.1101/227553

**Authors:** Brittany Jack, David M. Mueller, Ann C. Fee, Ashley Tetlow, Prachee Avasthi

## Abstract

Flagella of the unicellular green alga *Chlamydomonas reinhardtii* are nearly identical to cilia of mammalian cells and provide an excellent model to study ciliogenesis. These biflagellated cells have two actin genes: one encoding a conventional actin (IDA5) and the other encoding a divergent novel actin-like protein (NAP1). Previously, we described a role for actin in the regulation of flagella-building intraflagellar transport machinery. Here, we probe how actin redundancy contributes to this process using a *nap1* mutant *Chlamydomonas* strain. Disruption of a single actin allows normal or slower incorporation but complete flagellar assembly. However, when we disrupt both actins using Latrunculin B (LatB) treatment on the *nap1* mutant background, we find flagellar growth from newly synthesized limiting flagellar proteins is actin-dependent. Upon total actin disruption during flagellar assembly, transmission electron microscopy identified an accumulation of Golgi-adjacent vesicles, suggesting impaired vesicular trafficking may be the mechanism by which actin supports flagellar growth from new flagellar proteins. We also find there is a mislocalization of a key transition zone gating and ciliopathy protein, NPHP-4. Extended (2 hour) treatment with LatB, a condition under which NAP1 is upregulated, restores NPHP-4 localization. This suggests NAP1 can perform the functions of conventional actin at the transition zone. Our experiments demonstrate that each stage of flagellar biogenesis requires redundant actin function to varying degrees, with an absolute requirement for these actins in transport of Golgi-adjacent vesicles and flagellar incorporation of newly synthesized proteins.

## INTRODUCTION

Assembly and composition of the eukaryotic flagellum (also known as the cilium) are critical for signaling and development in most cell-types in the human body. The flagella of the green alga *Chlamydomonas reinhardtii* are essentially identical to the cilia of mammalian cells and provide an excellent model to study cell signaling, cell motility, and regulation of ciliary assembly. To date, the known mechanisms dictating the behavior of these organelles are largely dependent on the microtubule cytoskeleton. The flagellum is composed of microtubules that extend from a microtubule organizing center known as the basal body, and flagellar assembly requires control of microtubule dynamics. Trafficking from sites of cellular protein synthesis to their ultimate destination in flagella is also thought to occur on microtubule tracks (Tai et al., 1999).

However, evidence for the role of actin, another major cytoskeletal component, in ciliary regulation is increasing. In mammalian cells, disruption of actin leads to increases in both ciliary length and percentage of ciliated cells (Kohli et al., 2017; Sharma et al., 2011), which may be due to roles for actin in basal body migration, docking, and stabilization (Kim et al., 2010; Pan et al., 2007; Park et al., 2008; Tu et al., 2017; Yeyati et al., 2017). A recent study showed loss of Myosin-Va, an actin-based motor protein involved in trafficking of secretory vesicles from the post-Golgi to the plasma membrane, resulted in reduced ciliation. Disruption of Myosin-Va function stops the formation of the elongated ciliary membrane. This new result suggests actin and Myosin-Va are required for microtubule-dependent trafficking of preciliary and ciliary vesicles to the base of the cilia and therefore necessary for ciliogenesis (Tang, et. al, 2018). Actin is also required for vesicle budding in the endocytic pathway (Girao et al., 2008), which may influence ciliary assembly as there is a trafficking pathway connecting the endocytic compartments to a vesicular compartment involved in ciliary assembly (Kim et al., 2010). Membrane remodeling for both ciliary exocytosis (Nager et al., 2017) and ciliary resorption are actin dependent processes (Saito et al., 2017). In summary, the current understanding is that actin networks (that potentially block cortical access of basal bodies and ciliary proteins) inhibit cilium formation and elongation in mammalian cells but are also required for membrane trafficking to support ciliogenesis. Here, using an algal model system, we show a broader requirement for filamentous actin in flagellar protein synthesis, trafficking, and incorporation of proteins into an assembling flagellum.

Using *Chlamydomonas reinhardtii* as a model to interrogate flagellar dynamics, we are able to induce flagellar severing on demand to allow synchronous and successive rounds of flagellar regeneration. *Chlamydomonas* has two actin genes, one that encodes a conventional actin (IDA5) and another that encodes a novel actin-like protein (NAP1) with ~65% homology to mammalian actin (Hirono et al., 2003). *Chlamydomonas* cells treated with cytochalasin D, and actin polymerization inhibitor, exhibit flagellar shortening and regrowth upon washout of the drug. Suggesting a role for actin in flagellar maintenance (Dentler and Adams, 1992). *ida5* mutants and myosin inhibited cells had impaired flagellar motor recruitment to basal bodies, impaired entry of motors into the flagellar compartment, and ultimately a reduced initial flagellar growth rate (Avasthi et al., 2014). Despite defects in flagellar protein recruitment and flagellar assembly in *ida5* conventional actin mutants, these mutants ultimately grow flagella to wild-type lengths. Since *ida5* mutant flagella eventually reach wild type length, the degree to which actin is required is still in question.

In this study, we asked if more severe flagellar biogenesis phenotypes in IDA5 deficient cells are masked by redundant roles of the second *Chlamydomonas* actin NAP1, as *NAP1* expression increases upon IDA5 loss or disruption (Onishi et al., 2016). Using genetic and chemical perturbations of both *Chlamydomonas* actins, we find that the two actin genes have overlapping functions that include flagellar protein synthesis, vesicular trafficking, and composition of the flagellar gate.

## RESULTS

### Simultaneous disruption of all filamentous actins within *Chlamydomonas cells*

To investigate if *NAP1* contributes to the actin-dependent flagellar assembly functions previously identified for the conventional actin *IDA5* (Avasthi et al., 2014), we obtained a *nap1* null mutant strain, which was isolated based on its sensitivity to Latrunculin B (LatB), a conventional actin depolymerizing agent (Onishi et al., 2016). Here we use both a genetic and chemical approach to understand the role of actin in flagellar assembly (**Figure 1A**). LatB has been shown to disrupt filamentous actin in *Chlamydomonas reinhardtii (Avasthi et al., 2014; Onishi et al., 2016).* Treatment with LatB for ten minutes is sufficient to disrupt actin filaments in wild-type cells (Onishi et al., 2016). To confirm that LatB disrupts actin filaments under our experimental conditions, we fixed and stained the *nap1* mutant strain with phalloidin, a filamentous actin probe (**Figure 1**). We see actin filaments are present in the *nap1* mutants prior to drug treatment (**Figure 1B**). Previous attempts to visualize actin filaments with phalloidin have not been successful in vegetative *Chlamydomonas* cells. However, by selecting for cell health, considering the cell cycle, and shortening the incubation time to decrease background signal (Craig et al., in preparation), we were able to improve the signal to noise ratio and achieve specific filamentous actin labeling. Upon LatB treatment, these filaments disappear (**Figure 1D**). *Chlamydomonas* cells are known to upregulate NAP1 upon LatB treatment resulting in the return of filamentous signal, which represents the LatB-insensitive NAP1 population (Onishi 2016 and 2018). However, in *nap1* mutants, this upregulation cannot occur. Therefore, we do not see filaments return two hours post LatB treatment (**Figure 1D**). These results indicate that during the experiments presented here, we are testing phenotypes under conditions of no filamentous actin (neither IDA5 nor NAP1) in LatB-treated *nap1* mutant cells.

**Figure 1.**
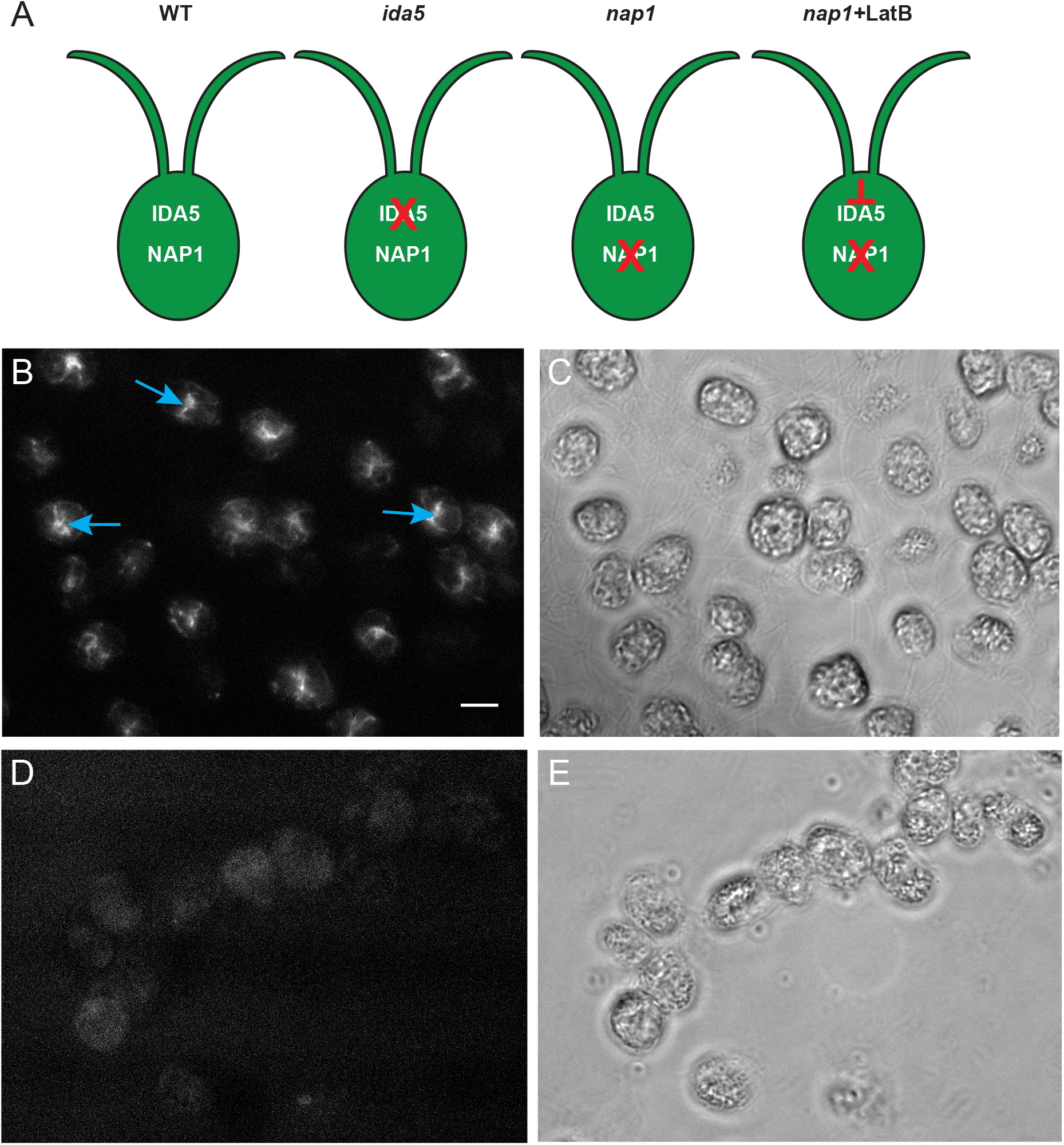
Latrunculin B (LatB) disrupts actin filaments on *nap1* mutant background. **A**. Genetic and chemical approach to investigate the role of actin in flagellar assembly. **B**. *nap1* mutant cells stained with phalloidin show filamentous actin (cyan arrows). **C**. Phase image of *nap1* mutant cells prior to LatB treatment. **D**. *nap1* cells treated with 10μM LatB and stained with phalloidin do not show filamentous actin. **E**. Phase image of *nap1* mutant cells after 2 hours of LatB treatment. Scale bar represents 5 μm.

### Flagellar length maintenance and assembly requires at least one functional actin

Cells with genetic disruption of *NAP1* and chemical perturbation of *IDA5* filaments lack all filamentous actins and ultimately cannot survive for long periods of time (Onishi et al., 2016). However, acute perturbation using LatB on the *nap1* mutant background allows us to probe the functions of both actins on the shorter time scales needed for assessing flagellar dynamics (**Figure S1**). When wild-type and *nap1* mutants are treated with 10μM LatB, wild-type flagella shorten but eventually recover while *nap1* mutant flagella continue to shorten (**Figure 2A**). To test if flagellar defects are due to indirect effects of actin disruption on microtubule organization, we labeled microtubules and confirmed that there were no gross defects in microtubule organization or microtubule number in LatB treated *nap1* mutants (**Figure S2**). These data suggest that some form of filamentous actin, either *IDA5* or *NAP1,* is required for flagellar maintenance.

**Figure 2.**
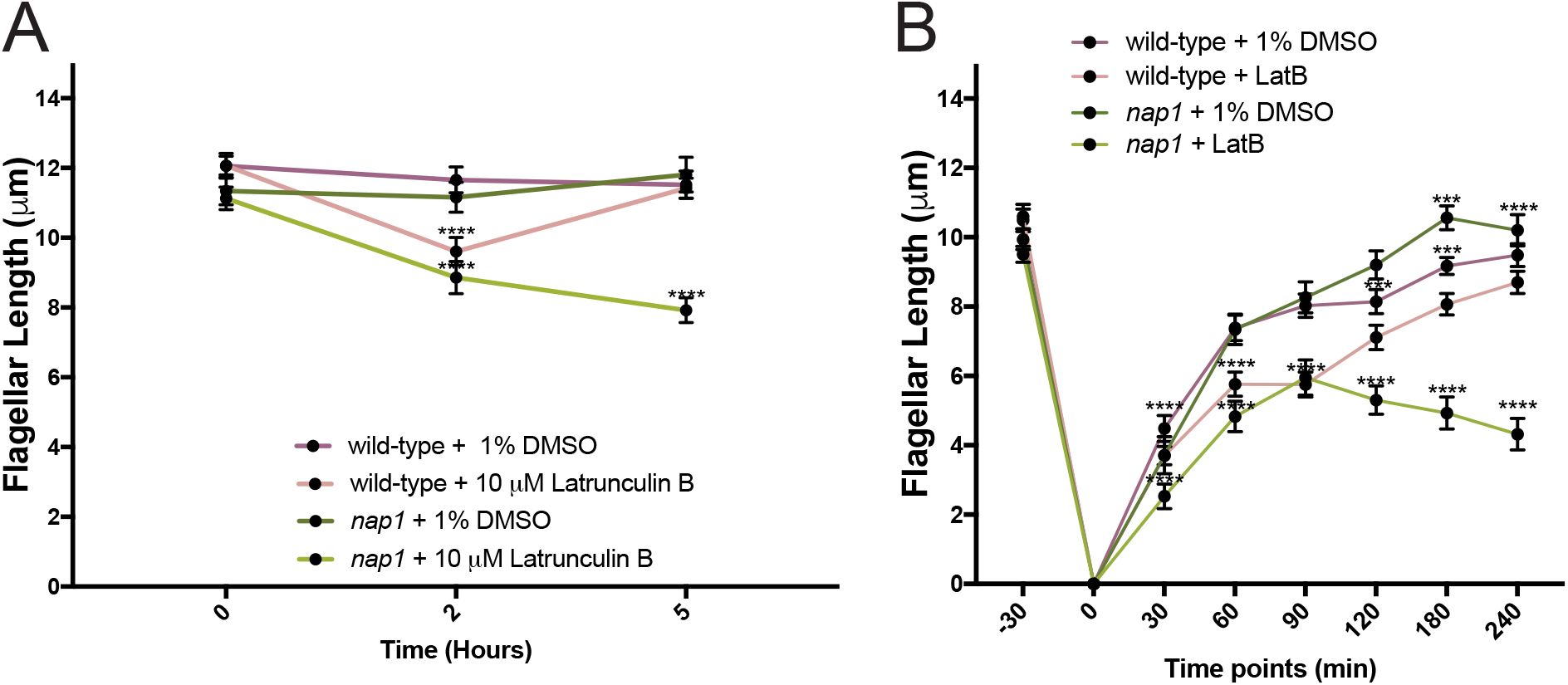
Actin filaments are necessary for flagella length maintenance and full flagellar assembly. **A**. Cells of each type were exposed to 10μM LatB. WT flagella shorten but recover while nap1 mutants + LatB cannot recover. Error bars represent 95% confidence interval (n=50). **B**. When all actin is disrupted in nap1 mutant cells we see the flagella only grow to half length. Flagellar lengths were assessed at various time points. Error bars represent 95% confidence interval. (***p<0.001, ****p<0.0001)

*ida5* mutants initially assemble their flagella more slowly but ultimately reach wild-type length (Avasthi et al., 2014). However, given that loss of all actins prevented flagellar maintenance (**Figure 2A**), we investigated if flagella could fully assemble when both actins are perturbed by deflagellating wild-type and *nap1* mutants via pH shock and regenerating flagella in the presence of LatB. When *IDA5* is disrupted on the *nap1* mutant background, flagella cannot grow beyond half-length of a typical flagellum (**Figure 2B**).

### Flagellar protein synthesis occurs but is reduced upon actin disruption

*Chlamydomonas* cells treated with the protein synthesis inhibitor cycloheximide following severing were previously shown to grow to half-length (Rosenbaum et. al, 1969). This half-length growth in cycloheximide utilizes already-synthesized flagellar proteins and demonstrates that synthesis of new limiting flagellar protein is required for full flagellar assembly. Given, actin’s known roles in transcription (Miralles and Visa, 2006), it may be involved in the deflagellation induced expression of flagellar proteins and failure to synthesize new flagellar protein may be one explanation for half-length growth in LatB-treated *nap1* mutants. To test this, we indirectly determined if actin loss blocked flagellar protein synthesis. Synthesis of new flagellar proteins can be quantified using cumulative flagellar growth beyond half-length (the amount of existing precursors) using an assay diagramed in **Figure 3A** and first described by Lefebvre et al. in 1978. To test actin’s effect on the amount of new protein synthesis at different time points following flagellar severing, *nap1* mutants are deflagellated and treated with 10μM LatB at 30 minute intervals to allow whatever protein synthesis and flagellar assembly that will take place under these conditions to proceed. Cells are deflagellated a second time, LatB is washed out, and all limiting flagellar proteins already synthesized are incorporated into flagella upon addition of cycloheximide. By comparing the cumulative length of flagella under actin perturbed conditions with half-length flagellar growth in cycloheximide (when zero protein synthesis takes place), we can infer the extent to which new limiting flagellar proteins were synthesized in the LatB treatment period (Jack and Avasthi, 2018). We see that loss of all actins does indeed reduce the amount of limiting flagellar proteins synthesized (**Figure 3B**) using this method in which flagellar length/growth is a proxy for limiting flagellar protein synthesis.

**Figure 3.**
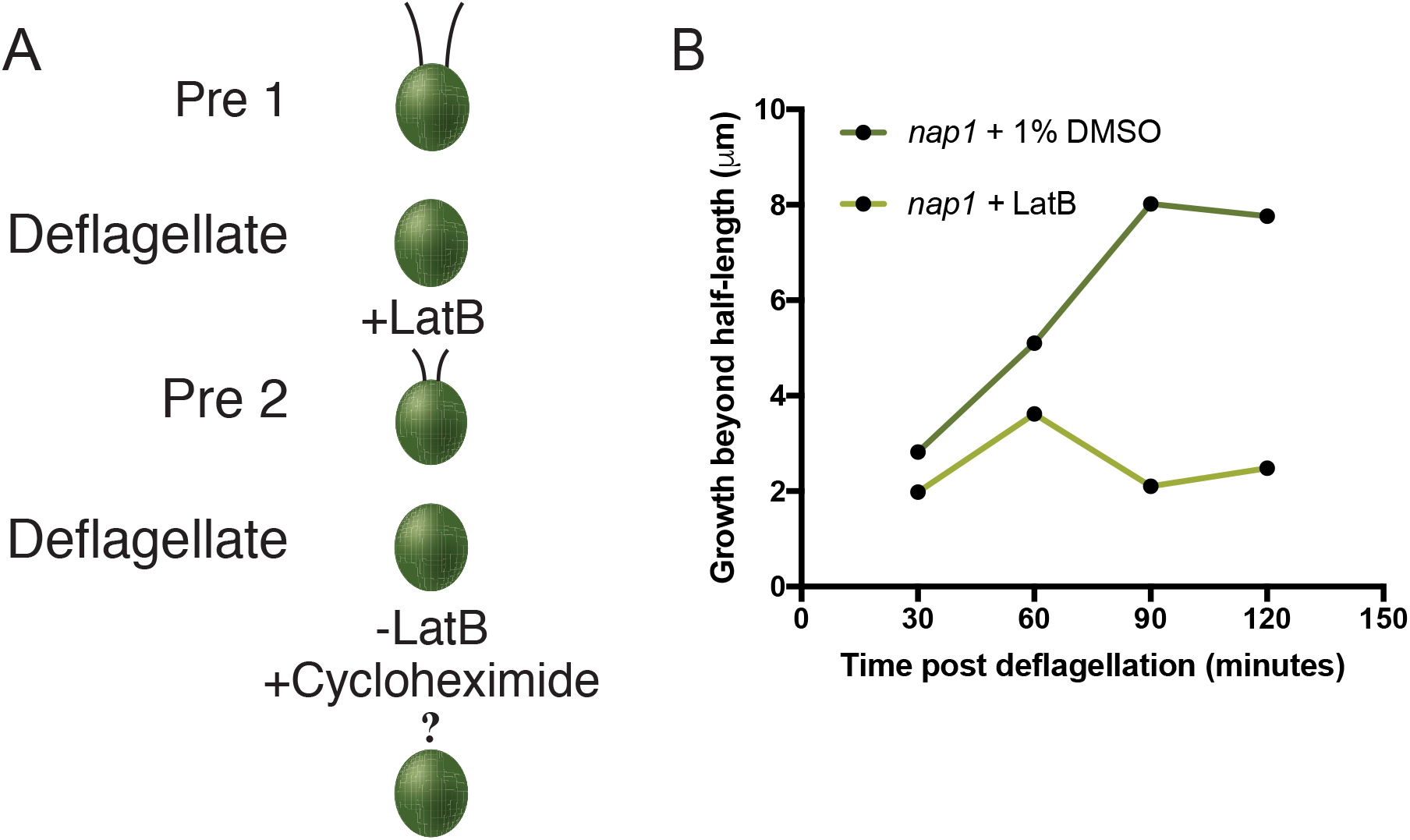
Flagellar protein synthesis is reduced when actins are disrupted. **A**. Schematic representation of new protein synthesis assay. **B**. Cells were deflagellated in the presence of 10μM LatB and deflagellated a second time washing out the LatB and adding in 10μg/mL Cycloheximide. This process of double deflagellation allows for quantification of newly synthesized limiting flagellar protein when actins are disrupted. Fewer new flagellar proteins are synthesized upon disruption of both actins. For all measurements (n=30).

### Dual actin disruption blocks all flagellar growth when flagellar precursor proteins are depleted

If limiting flagellar proteins can be synthesized under actin depleted conditions the question remains as to why these flagella cannot assemble beyond half length. We asked whether the flagellar growth represents incorporation of only the existing pool of cytoplasmic flagellar proteins but not newly synthesized flagellar proteins. To test this, we depleted the limiting flagellar proteins in the cytoplasmic pool prior to flagellar assembly under actin disrupting conditions. A schematic of this experiment is shown in **Figure 4A**. In this precursor protein-depleted condition, the cell must synthesize proteins de novo, traffic and incorporate them into the flagella. In this context, complete disruption of actin blocks all flagellar growth (**Figure 4B**). This suggests that 1) the flagella can reach half-length in LatB treated *nap1* mutants exclusively from incorporation of the existing pool of flagellar proteins, and 2) that some form of actin is an absolute requirement for newly synthesized proteins to be incorporated into growing flagella.

**Figure 4.**
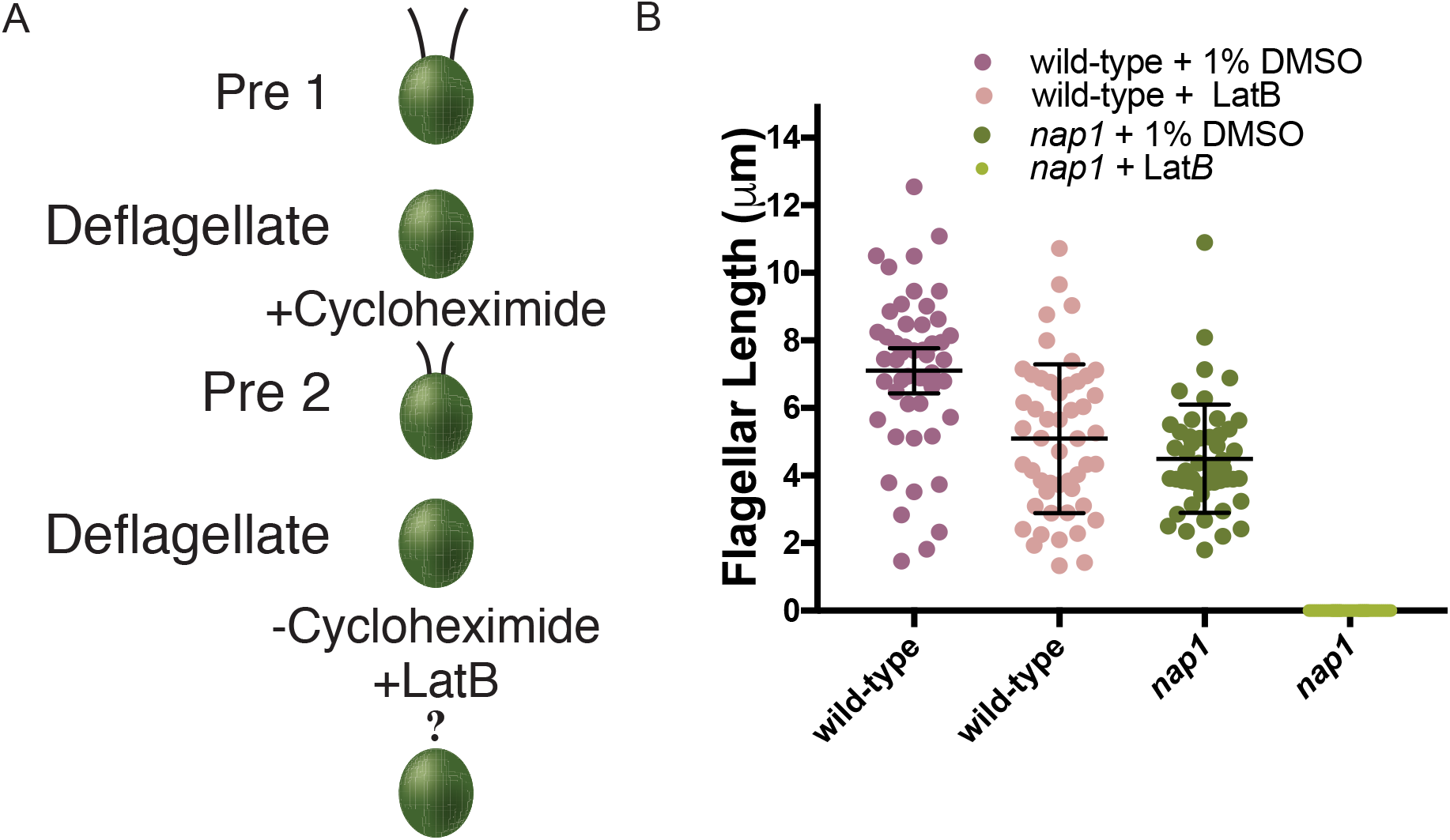
Functional actin is necessary for incorporation of newly synthesized flagellar protein. **A**. Schematic of new flagellar protein incorporation assay. **B**. New flagellar protein incorporation assay, measuring the effects of Lat B on *nap1* mutant flagellar regeneration under precursor pool depleting conditions. There is no flagellar growth using newly synthesized protein in *nap1* mutants when IDA5 actin is disrupted. Error bars are 95% confidence interval (n=50).

### Golgi-adjacent vesicles accumulate upon filament actin disruption

To assemble a flagellum, all the necessary components must arrive at the base of the cilia as there are no ribosomes within the ciliary compartment (Deiner et. al, 2015). We have shown there is no flagellar growth under conditions where components needed to build a flagellum must be newly synthesized, trafficked and incorporated (**Figure 4**). The question remains as to why this newly synthesized protein (**Figure 3**) cannot be incorporated into a flagellum. To examine if there are gross abnormalities in Golgi morphology which could possibly be the cause of this incorporation problem, *nap1* mutant flagella were severed to induce conditions that require additional protein synthesis and trafficking. These cells were visualized by transmission electron microscopy (TEM) and we find no noticeable abnormalities in Golgi cisternae in control cells (**Figure 5A and B**, magenta arrows). However, under actin deficient conditions, we see a dramatic accumulation of Golgi-adjacent vesicles (**Figure 5C and D, magenta arrows**). To determine the extent of the accumulation and identify any additional defects, we quantified the number (**Figure 5E**) and size (**Figure 5F**) of these vesicles. The four analyzed control and actin-deficient cells have 14 and 43 total vesicles respectively while their diameter is unchanged.

**Figure 5.**
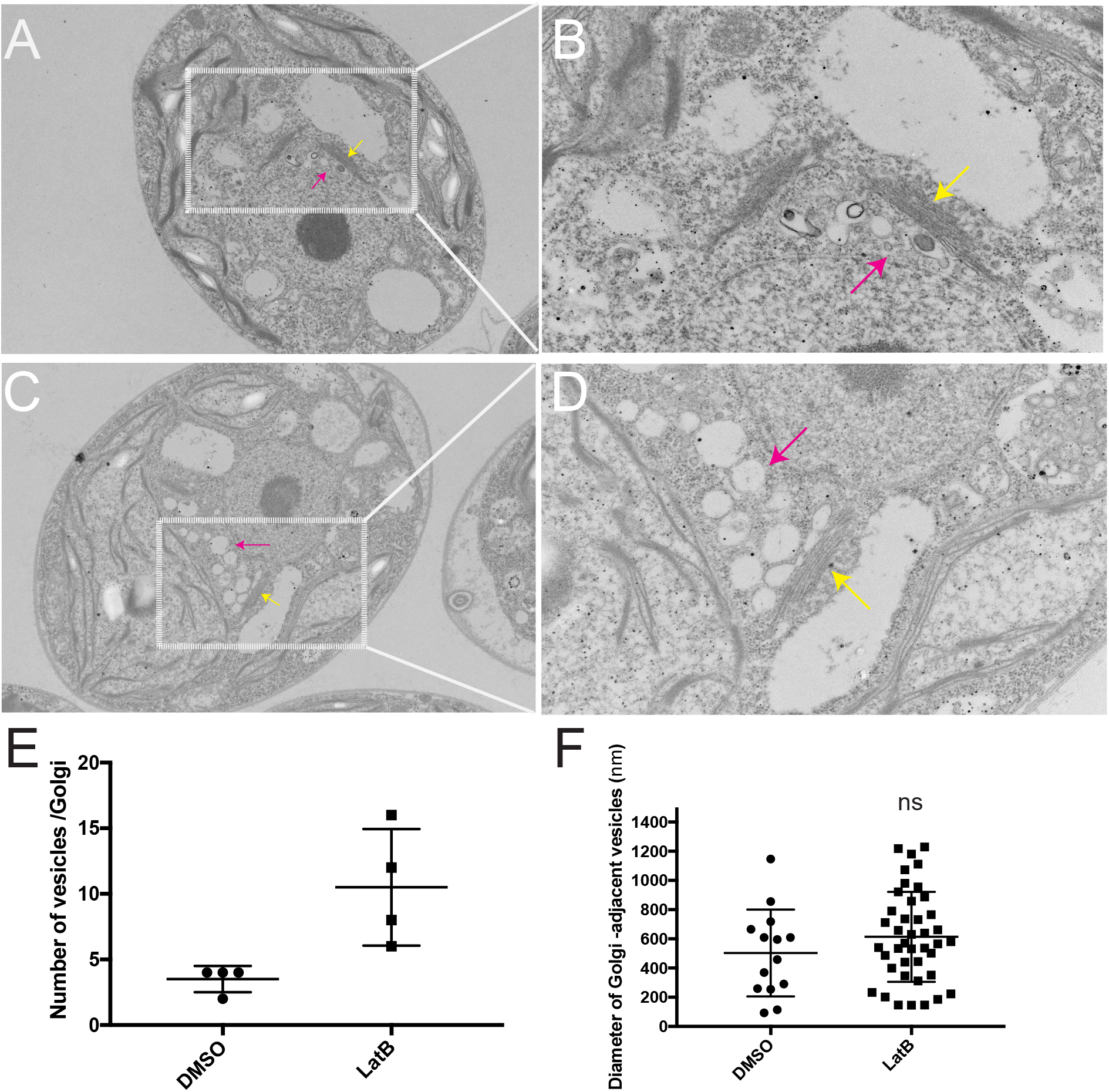
Actin depletion results in accumulation of Golgi-adjacent vesicles upon flagellar regeneration. **A**. *nap1* cells treated with DMSO 30 min post-deflagellation at 4000X magnification. **B**. *nap1* cells treated with DMSO 30 min post-deflagellation at 10000X magnification. This is an independent image of the area outlined in panel A taken at a higher magnification. **C**. *nap1* cells treated with LatB 30 min post-deflagellation at 4000X magnification. **D**. *nap1* cells treated with LatB 30 min post-deflagellation. Independent image of area outlined in panel C taken at 10000X magnification. **Yellow** arrows point to the Golgi apparatus and **magenta** arrows pointing to Golgi-adjacent vesicles. **E**. Quantification of number of vesicles per Golgi (n=4). **F**. Quantification of diameter of Golgi-adjacent vesicles. Difference is not significant.

### Actin loss produces flagellar gating defects

During regeneration in LatB treated *nap1* mutants (**Figure 2B**), we noticed that the flagellar growth to half-length appeared slower than cells containing both actins. To test whether or not this represents a defect in incorporating proteins from the existing pool of flagellar precursors, we simultaneously treated *nap1* mutants with cycloheximide and LatB following deflagellation. LatB treated cells (both wild-type cells that take time to upregulate *NAP1* and *nap1* mutants) incorporate the existing pool of flagellar precursors at a slower rate than cells with intact actins (**Figure 6F**). We also previously found that during flagellar assembly, there was a small range of flagellar lengths (7-9μm) in which flagellar motor recruitment to basal bodies in *ida5* mutants was comparable to controls but motor entry into flagella was reduced (Avasthi et al., 2014). These data in conjunction with slow incorporation of existing precursors, suggest a role for actin in gating of material already accumulated at the flagellar base. A region thought to be critical for gating functions is the transition zone at the base of flagella which has connectors between the microtubule core and flagellar membrane. The transition zone also houses proteins found to regulate the composition of flagella (Awata et al., 2014). Actin itself is found in the transition zone of flagella (Diener DR, 2015) and may function to transport or anchor the transition zone gating proteins. *NPHP-4,* a gene mutated in the cilium-related kidney disease nephronophthisis, is a crucial component of the ciliary gate that controls entry of both membrane-associated and soluble proteins (Awata et al., 2014). We used a strain expressing HA-tagged NPHP-4 to test the effects of functional actin loss on NPHP-4 localization. In control cells stained with anti-HA antibody, NPHP-4 localizes at two apical spots at the base of flagella (**Figure 6A, cyan arrows**). In cells treated with LatB for 10 and 30 minutes, we saw a dramatic loss of apical NPHP-4 localization (**Figure 6 B and C, magenta arrows**). Occasionally we could see some remaining apical staining and the results are quantified in **Figure 6 E**. These results demonstrate that acute disruption of actin with LatB treatment (at a time point prior to *NAP1* upregulation) causes a reorganization of the transition zone for loss of a known transition zone gating protein, NPHP-4. Extended treatment of LatB (2 hours), a condition in which NAP1 is upregulated, shows a recovery of NPHP-4 to the apical part of the cell. (**Figure 6D**, **cyan arrows and E, gray bars**). This suggests NAP1 can perform this function of conventional actin at the transition zone.

**Figure 6.**
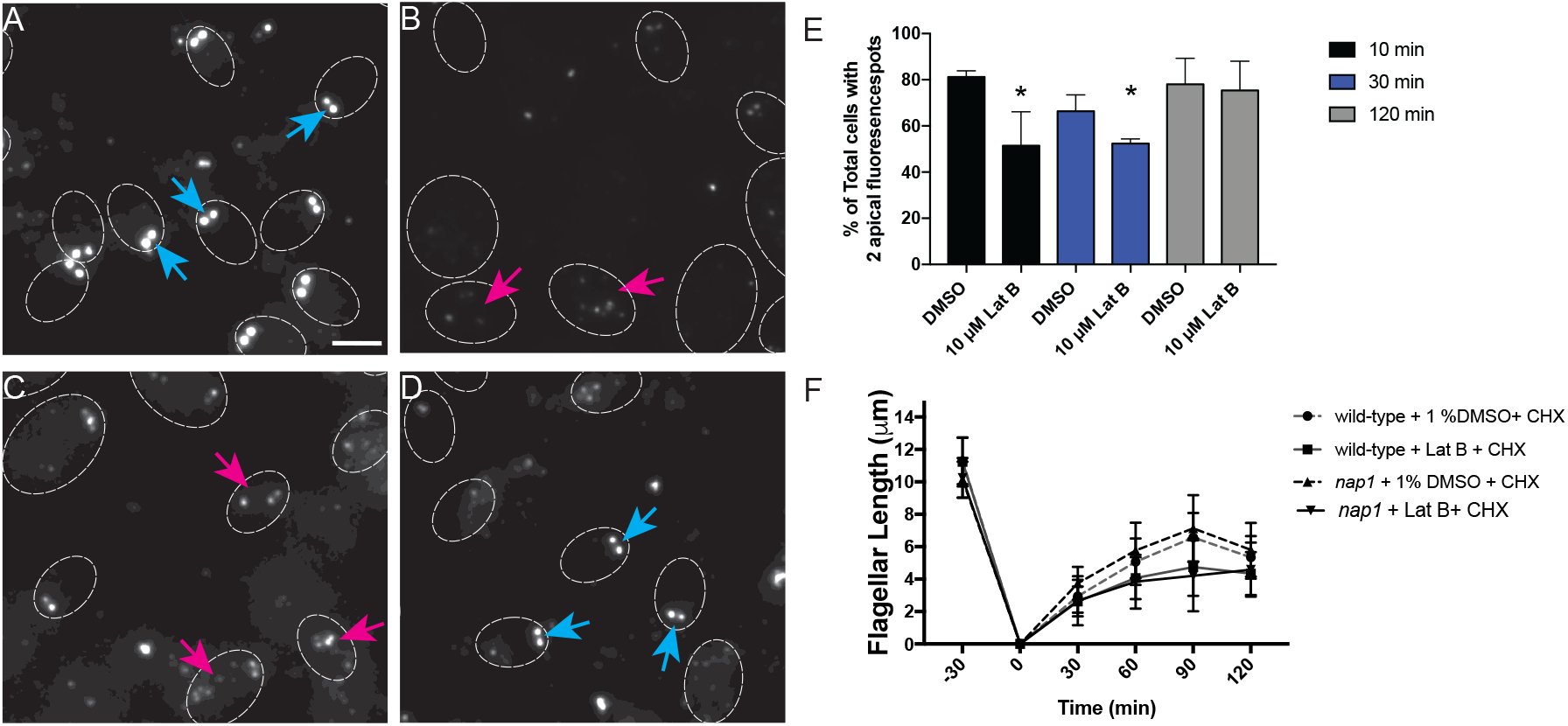
The transition zone gating protein NPHP-4 and incorporation of flagellar proteins are affected upon disruption of Chlamydomonas actins. **A**. Cells expressing HA tagged NPHP-4 were stained with HA primary antibody and localization is seen at the base of the flagella (cyan arrows). **B**. Cells treated with 10 μM LatB for 10 min were stained with the HA antibody. We see mislocalization of the NPHP4 protein (magenta arrows). **C**. Cells treated with 10 μM LatB for 30 min were stained with the HA antibody. **D**. Cells treated with 10 μM LatB for 120 min were stained with the HA antibody. Cyan arrows indicate cells with a recovery of apical NPHP4 localization. **E**. Percentage of cells with two apical fluorescence spots upon NPHP-4 labeling are quantified (*:p<0.05, N=3). **F**. Cycloheximide is used to stop new protein synthesis. The addition of LatB (or 1% DMSO control) is to test the effect actin disruption has on incorporating flagellar protein. Flagellar lengths were assessed every 30 minutes for 2 hours. Error bars represent 95 % confidence. Scale bar represents 5 μm.

## DISCUSSION

In this study, we acutely disrupt all filamentous actin in *Chlamydomonas* through treatment of *nap1* mutants with LatB (**Figure 1**). Recent transcriptomic analyses show that depolymerization of F-actin by LatB treatment in these cells induces an upregulation not only of NAP1, but several hundred other genes including a general upregulation of the ubiquitin proteasome system to monitor actin filament integrity and prevent dominant negative effects on NAP1 by monomeric IDA5 (Onishi et al., 2018; Onishi et al., 2016). Loss of IDA5 in the *nap1* mutant through LatB treatment does not affect which genes are upregulated (Onishi et al., 2018). The upregulation of these genes may influence the phenotypes seen in this study and must be considered as potential contributing factors.

Our data supports a requirement of at least one form of *Chlamydomonas* actin for flagellar length maintenance and full flagellar assembly. Anterograde flagellar motor complexes, which are required for assembly of the flagellum, utilize newly recruited proteins from the cell body pool (Wingfield et al., 2017). With constant turnover and a demand for continuous recruitment to the base of flagella it appears that actin plays a role in this recruitment (Avasthi et al., 2014). We find functional actin is required for normal flagellar protein synthesis and normal incorporation of existing proteins (**Figure 7**). Importantly, we found that actin-deficient cells cannot grow flagella at all when new flagellar proteins must be 1) synthesized, 2) trafficked, and 3) incorporated (**Figure 4**). However, some level of flagellar protein synthesis occurs (given the reduced but non-zero values for LatB treated *nap1* mutants in **Figure 3B**). Together, these data support a model in which actin is an absolute requirement for entry of proteins into a usable cytoplasmic pool.

**Figure 7.**
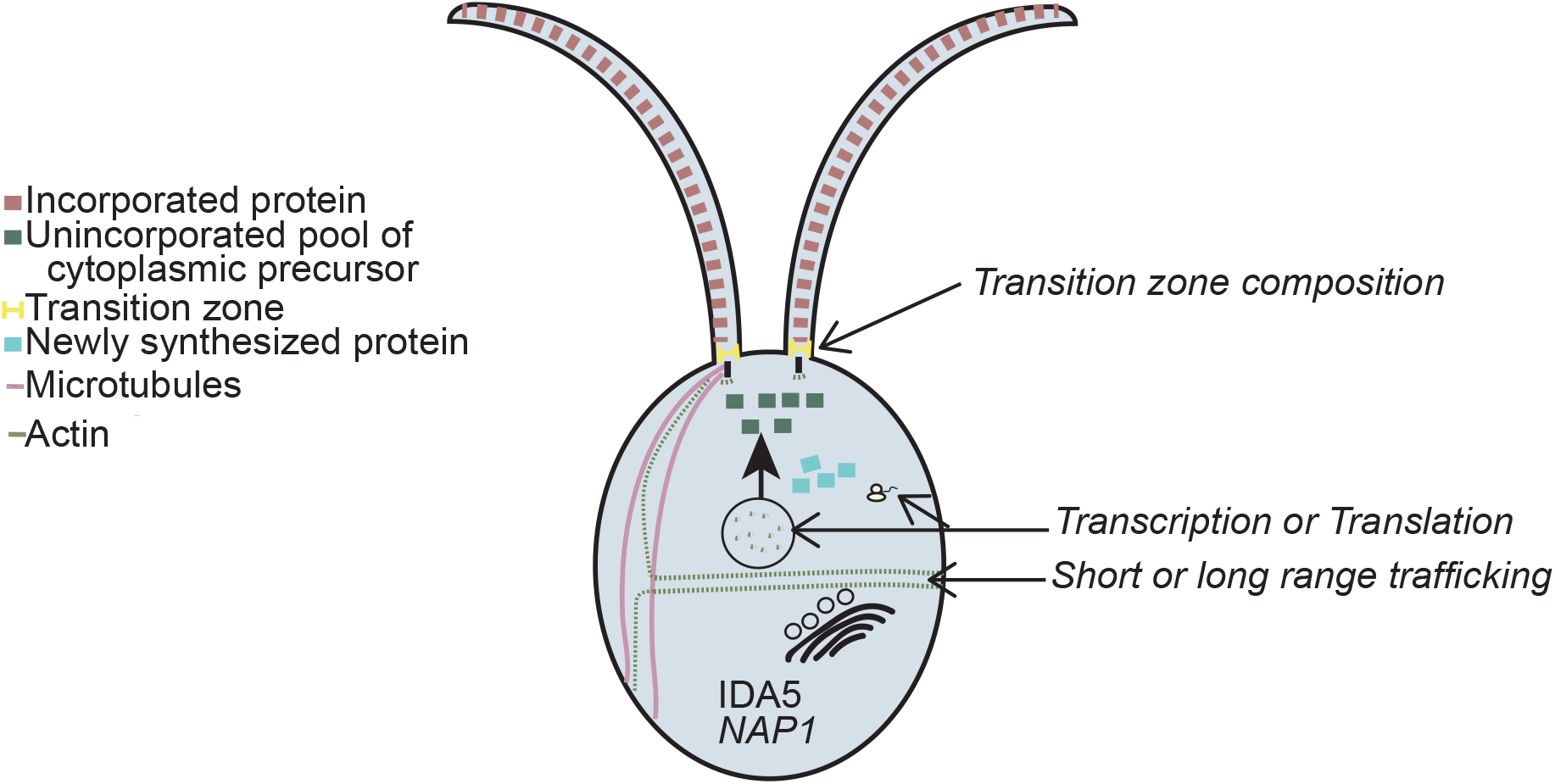
Model of actin dependent ciliary functions. Loss of all functional actins via depolymerization of IDA5 actin in nap1 mutants results in no flagellar growth using newly synthesized protein. Disruption of both actins causes slower protein incorporation and less newly synthesized flagellar protein but cannot account for the inability to grow flagella. We hypothesize functional actin is required for short of long range trafficking of newly synthesized protein to the flagella base.

Flagellar membrane proteins/vesicles originate from the Golgi (Nachury et al., 2010; Rohatgi and Snell, 2010). Further, in *Chlamydomonas,* collapse of the Golgi upon treatment with Brefedlin A results in a shortening of the flagella, suggesting that limiting protein or membrane destined for the flagella are being exported from the Golgi (Dentler, 2013). Therefore, we reasoned that in the absence of actin, flagellar proteins or membrane cannot be properly recruited, sorted, or released from the Golgi. Our TEM results show that Golgi morphology is not dramatically altered but there is an accumulation of what appear to be secretory vesicles adjacent to the *trans* face of the Golgi (the *trans* face identified by narrower cisternae (Engel et al., 2015) in close proximity to larger secretory vesicles). Given the *trans* Golgi vesicle accumulation and previous studies implicating a role of myosin in the trafficking of flagellum-directed cargo (Avasthi, et al, 2014) we propose a model in which acto-myosin dependent transport is required for either short or long range trafficking of flagellum-bound vesicles after protein synthesis. Further studies evaluating the role of actin in flagellar protein sorting and trafficking, including the role of individual myosins that have been implicated by us (Avasthi et al., 2014) and others (Assis et al., 2017), are currently underway.

Also novel is the finding that actin is involved in flagellar protein synthesis (**Figure 3**). While actins are known to localize to the nucleus (Pederson and Aebi, 2002) and interact with transcription factors, chromatin remodeling complexes, and RNA polymerases (Miralles and Visa, 2006), it is not well defined whether all or a subset of transcripts require actin function. Given the extensive interaction of actin with core transcriptional machinery, perhaps it is more unexpected that some flagellar protein synthesis can indeed occur in the absence of functional actins. The identity of the proteins that are reduced under these conditions is still unknown. Also, while we have taken two different approaches to test protein synthesis in *Chlamydomonas* under actin depleted conditions, it remains to be determined via proteomics and transcriptomics whether our assays reflect a decrease in translation or transcription.

Lastly, we found a significant disruption of NPHP-4 transition zone localization (**Figure 5B and D**). In *Chlamydomonas,* actin was identified through biochemical purification as a transition zone protein and may serve as a scaffold in the region. While NPHP-4 turnover appears more static in *Chlamydomonas* cells (Awata et al., 2014), actin and myosins found in the peri-basal body region (Assis et al., 2017; Avasthi et al., 2014) may be involved in turnover of transition zone components in mammalian cells where the NPHP-4 turnover rate is high (Takao et al., 2017). *Nphp-4* mutants also show decreased membrane protein entry into flagella, suggesting the protein is important in regulating flagellar composition. Membrane associated proteins in *C. elegans* (Williams et al., 2011) and soluble housekeeping proteins in *Chlamydomonas* (Awata et al., 2014) also inappropriately accumulate within cilia in *nphp-4* mutants, suggesting that there is a more general dysregulation of ciliary gating upon NPHP-4 loss. Finally, *nphp-4* mutants in *C. elegans* have ultrastructural abnormalities (Jauregui et al., 2008; Lambacher et al., 2016). Therefore, at minimum, actin disruption, which significantly affects NPHP-4 localization, is expected to encompass the full range of ciliary phenotypes associated with NPHP-4 loss.

*Chlamydomonas Nphp-4* mutants exhibit slow flagellar assembly that ultimately reach wild-type flagellar lengths (Awata et al., 2014), similar to *ida5* mutants (Avasthi et al., 2014). Is NPHP-4 loss then the primary cause of slow initial flagellar assembly in *ida5* mutants? We found that NPHP-4 reappears at the basal bodies after extended treatment of LatB on *nap1* mutant cells. These data suggest NAP1 is able to compensate for the functions of IDA5 at the transition zone. Although NAP1and IDA5 have some redundant functions, we still see slower rates of assembly in the *ida5* mutant, in which NAP1 is expressed and further upregulated during regeneration. This suggests either that NAP1 expression levels remain insufficient early on in regeneration to support proper NPHP-4 localization or that other factors outside of transition zone disruption are responsible for early regeneration defects in actin-disrupted cells.

Previous work (Avasthi et al., 2014) and the data presented here allow us to isolate specific steps of flagellar assembly that require normal actin dynamics. Actin has many cellular functions, including in organelle morphology, transcription, membrane dynamics, and polarized trafficking. We are now beginning to see how these conserved actin functions can dramatically influence flagellar biogenesis, a process previously thought to be controlled primarily through microtubule regulation.

## Supporting information

## ACKNOWLEDGEMENTS

We would like to thank Masayuki Onishi, John Pringle, and Fred Cross for providing the *nap1* mutant. Thank you to members of the Avasthi Lab for troubleshooting and manuscript feedback. Thank you to Evan Craig, whose optimized phalloidin staining protocol was used in this paper. Thanks to Bill Dentler for help with electron microscopy protocols and critical reading of a version of the manuscript. This work was funded by P20GM104936 and R35GM128702 (PA) and NSF GRFP 1518767 (BJ).

## MATERIALS AND METHODS

### Compounds

Latrunculin B, DMSO and Cycloheximide were purchased from Sigma and used at indicated concentrations and incubated for specified times.

### Strains

Wild-type, *ida5* mutants NPHP4-HAC were obtained from the *Chlamydomonas* stock center as CC-125 mt +, CC-3420, and CC-5115. The *nap1* mutant was a generous gift from Fred Cross, Masayuki Onishi, and John Pringle.

### Inhibitor Treatment and Flagellar Length Measurements

All strains were grown in liquid TAP medium for 24 hours prior to incubation with 10 μM Latrunculin B or 10 μg/mL for indicated times. Cells were fixed in 1% glutaraldehyde and imaged by DIC microscopy at 40 X magnification. Flagellar lengths were measured using ImageJ software by line segment tracing and spline fitting.

### Phalloidin Staining

Cells were grown in liquid TAP media 24 hours prior to experiment. Cells were incubated with either 10 μM LatB or 1% DMSO for 2 hours min. Cells were fixed to coverslips with 4% paraformaldehyde fixation and permeabilize with acetone. Cells were stained with phalloidin for 15 minutes (optimized time for bright signal) and washed with PBS prior to mounting.

### Flagellar Regeneration

Flagellar regeneration was induced by deflagellating cells via pH shock by adding 60μL of 0.5N acetic acid followed by 70 μL of 0.5N KOH to 1 mL of cells in liquid TAP. Cells were fixed with 1% glutaraldehyde at 0,30,60,90,120, and 240 minutes. Flagella were measured as described in the method above. Double deflagellation experiments for new protein incorporation were performed as described above (Figure 2A). The cells were deflagellated via pH shock and treated with 10μg/mL of Cycloheximide for 2 hours. The cells were deflagellated a second time and the cycloheximide was washed out and 10μM LatB was added. The cells were treated for 4 hours and samples were taken every 30 minutes. Flagella lengths were measured using ImageJ software by line segment tracing and spline fitting.

### Immunofluorescence Microscopy of NPHP4-HAC

Cells were grown in liquid TAP media 24 hours prior to experiment. Cells were incubated with either 10 μM LatB or 1% DMSO for 30 min. Cells were fixed to coverslips with 4% paraformaldehyde fixation and stained with primary HA anti-rat purchased from Sigma at (1:1000) dilution. The secondary antibody used was also purchased from Sigma and used at (1:1000) dilution as well. Slides were visualized on Nikon TiS microscope on the FITC channel.

### New protein synthesis quantification

To determine if the cell can synthesize protein under actin depleted conditions, we performed a double deflagellation experiment to quantify the amount of newly synthesized protein. The method is described in Figure 3A but briefly, the initial deflagellation in this experiment follows treatment with 10 μM LatB to allow protein synthesis under actin depleted conditions. The first deflagellation runs for a total of 2 hours but cells are deflagellated a second time every 30 minutes where the LatB is washed out and 10μg/mL cycloheximide is added in to prevent any new protein synthesis. The addition of cycloheximide allows only protein that was synthesized during LatB treatment to be incorporated into the assembling flagellum. Growth beyond half-length is quantified as newly synthesized protein.

### Electron Microscopy

For thin sections, cells were deflagellated via pH shock by adding 60μL of 0.5N acetic acid followed by 70 μL of 0.5N KOH to 1 mL of cells in liquid TAP. Cells regenerated flagella for 30 min and were then fixed in 2% glutaraldehyde for 15-20 min at room temp. The cells were gently pelleted using 1X G and then fixed for 1 hr at room temp and overnight at 4 degrees. The staining protocol was followed according to Dentler and Adams, 1992.

