## Supplementary material for "Partially redundant actin genes in *Chlamydomonas* control flagellum-directed traffic and transition zone organization"

### Supplemental Material

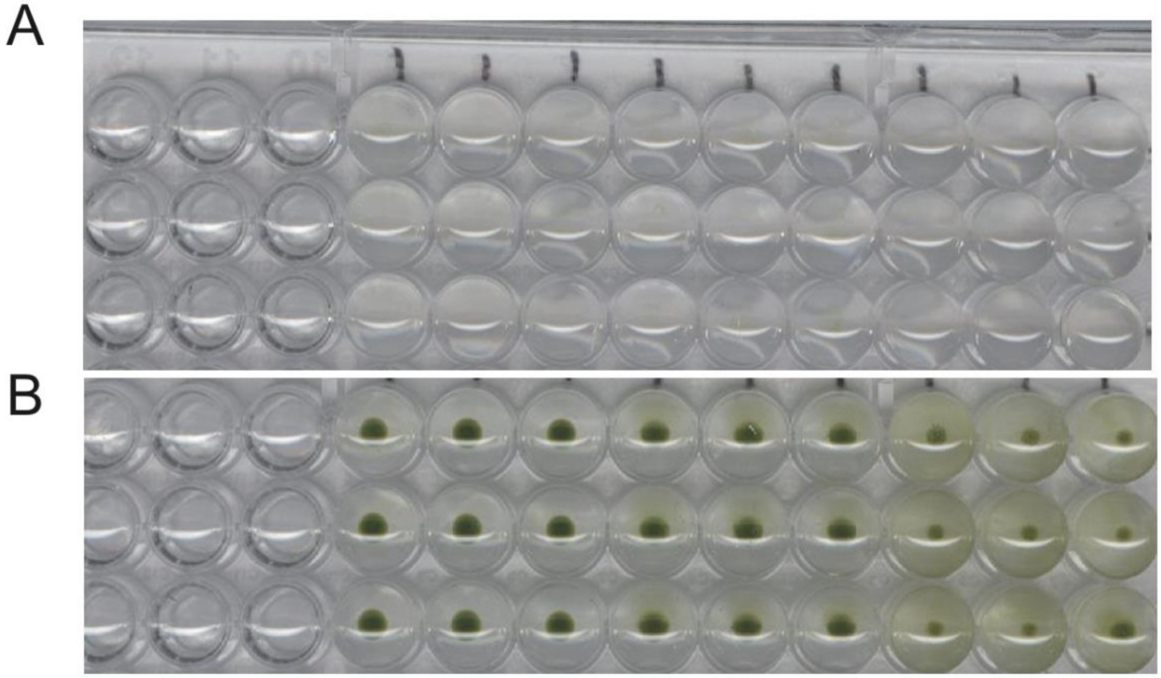

**Supplemental Figure 1. Chlamydomonas strains survive LatB treatment for at least up to 5 hours Related to Figure 2. A.** wild-type, ida5, and nap1 strains treated with 10µM Lat B for 5 hours. LatB was washed out and an aliquot of the cells were added to 99 µL of liquid TAP media immediately following experiment. **B**. 5 days after re-suspension in TAP media growth was assessed and we see growth of all strains indicating wild-type, ida5, and nap1 can survive LatB treatment for at least 5 hours.

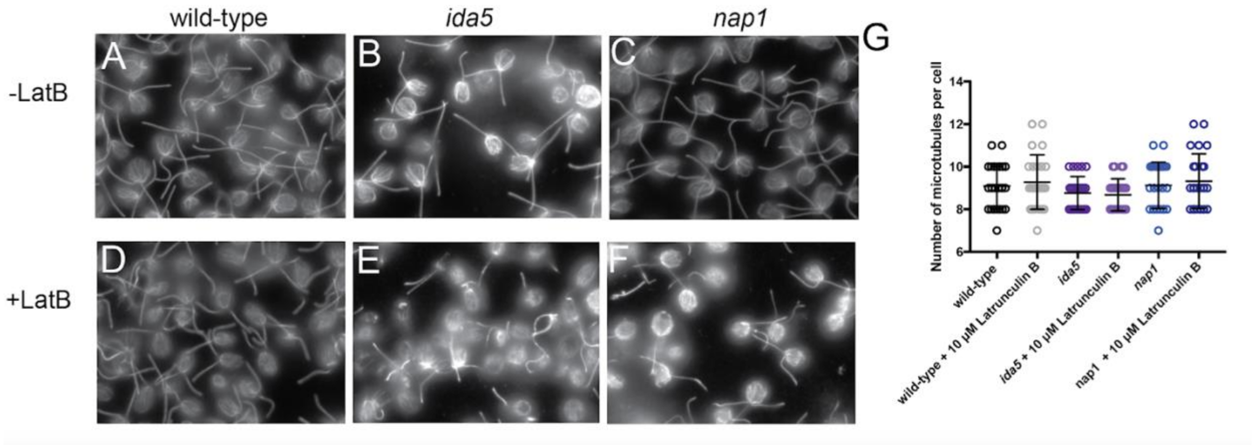


**Supplemental Figure 2. Actin disruption does not cause alteration in microtubule number Related to Figure 2. A.** Wild-type cells without LatB treatment. **B**. *ida5* mutants without LatB treatment. **C**. *nap1* mutants cells without LatB treatment. **D**. Wild-type cells with LatB treatment show no difference in number of microtubules. **E**. *ida5* mutants cells with LatB treatment show no difference in microtubule number. **F**. *nap1* mutant cells with LatB treatment show no difference in LatB microtubule number. **G**. Quantification of number of microtubules per cell.

Supplemental Experimental Procedures

**Cell Survival in LatB**

Wild-type, *ida5* mutants, and *nap1* mutants were grown in liquid culture 24 hours and underwent the double deflagellation experiment for new protein incorporation. Following the experiment 1 µL of cells were added to 99 µL of liquid TAP media in a 96 well plate. An image was taken immediately after the experiment and 5 days following to assess for growth. Growth of all strains indicate these cells can survive LatB treatment for at least 5 hours.

**Microtubule Staining**

Cells were grown in TAP medium 24 hours prior to treatment. The cells were treated with 10 µM Lat B for 30 minutes. The cells were then extracted into Eppendorf tubes and centrifuged for 2 minutes at 7500 rcf. The supernatant was removed and replaced with an equal amount of MT buffer. MT Buffer :30 mM HEPES (pH 7.2), 3mM EGTA, 1mM MgSO4, and 25 mM KCl in ddH20. Cells were adhered to a poly-Lysine-treated coverslip for five minutes. The coverslips were then fixed 4%paraformaldehyde, diluted in MT buffer, for five minutes. Next, the coverslips were treated with 200µL of 0.5% NP40, also diluted in MT buffer. The cells underwent methanol extraction at -20°C for 5 minutes. The coverslips were placed in a humidified chamber and treated with 100µL of 100% block (5% BSA and 1% cold water fish gelatin in 1X PBS) for 30 minutes at room temperature. Solution was tilted off and 100µL of 10% Normal Goat Serum (diluted in 100% block) were applied, the coverslips sat for 30 minutes at room temperature, in the humidified chamber. The cells received a solution of primary antibody (monoclonal anti-mouse alpha tubulin1:100 purchased from Sigma) diluted in 20% block. The cells received primary antibody treatment in the humidified chamber overnight at -4°C. The following day, the cells were washed three times, for ten minutes each time, in 1X PBS, then treated for thirty minutes with secondary antibody (1:1000 goat-anti-mouse-Alexa 488) at room temperature in the humidified chamber. Again, the coverslips cells were washed three times, for ten minutes each time, in 1X PBS. Finally, the coverslips were mounted on glass slides with 7µL of Fluoromount-G and imaged in the FITC channel, taking 0.2µM step Z-stack images on a Nikon TiS microscope.
